# Cortico-basal ganglia dynamics of global and selective response inhibition in humans

**DOI:** 10.64898/2026.01.20.700500

**Authors:** Cheol Soh, Mario Hervault, Nathan H. Chalkley, Kien Huynh, Qiang Zhang, Ergun Y. Uc, Jeremy D. W. Greenlee, Jan R. Wessel

**Affiliations:** Department of Psychological and Brain Sciences, University of Iowa, USA; Department of Neurology, University of Iowa Hospitals and Clinics, USA; Cognitive Control Collaborative, University of Iowa, USA; University of Grenoble, Grenoble, France; Neurology Service, Iowa City VA Medical Center, Iowa City, USA; Department of Neurosurgery, University of Iowa Hospitals and Clinics, USA

**Keywords:** Inhibitory Control, Response Inhibition, Subthalamic Nucleus, β bursts, Stop-Signal Task

## Abstract

Response inhibition is an important cognitive control mechanism, which enables flexible behavior by stopping inappropriate actions. Intracranial recordings across species have identified a neural circuit that implements response inhibition via the subthalamic nucleus of the basal ganglia. However, this work has been limited to simple tasks, in which unequivocal, salient “stop”-signals require the inhibition of all ongoing responses. Notably, response inhibition in the real world is substantially different. Real-world response inhibition is selective: it occurs only after specific salient signals (‘stimulus-selectivity’) and stops only specific movements, while others continue (‘response-selectivity’). If and how the fronto-subthalamic system implements selective inhibition is largely unknown. Here, we recorded subthalamic local field potentials and scalp-EEG in humans performing a novel, selective inhibition task. Salient signals either required stopping all initiated responses (global inhibition), stopping only some responses (response-selective inhibition), or continuing all responses – i.e., ignoring the signal (which ensures stimulus-selectivity). All three signals initially triggered a common fronto-subthalamic inhibitory process, signified by a rapid increase in β-burst activity. During global inhibition, subthalamic β-bursting subsequently increased above baseline, persisting for over a second. During response-selective inhibition, this activity was delayed, which enabled a second bout of disinhibition and allowed appropriate responses to continue. Throughout this period, frontal cortical and subthalamic β-band activity was tightly coupled. This shows that selective inhibition is implemented through rapid, context-dependent engagement and release of fronto-subthalamic inhibition. Moreover, subthalamic activity lasted substantially longer than assumed by classic behavioral-computational models. This supports recent theoretical models that assume protracted response inhibition during action-stopping.

## Introduction

Response inhibition is one of the fundamental cognitive control functions that enables flexible behavior (Verbruggen & Logan, 2009). It allows humans, non-human animals, and other cognitive agents to rapidly cancel inappropriate or obsolete actions (Chambers et al., 2009; Schmidt & Berke, 2017). Response inhibition is implemented by a fronto-basal ganglia neural circuit (Wiecki & Frank, 2013; Aron et al., 2014; Wessel, 2025). Frontal cortical regions signal the need to stop ongoing actions to the basal ganglia via two net-inhibitory motor pathways (indirect and hyperdirect, Jahanshahi et al., 2015; Chen et al., 2020; Wessel & Anderson, 2024). These pathways converge on the subthalamic nucleus (STN), which net-inhibits the thalamocortical motor system via broad projections to the output nuclei of the basal ganglia (Mink, 1996; Frank, 2006).

The vast majority of our knowledge about motor inhibition comes from stop-signal tasks (Logan & Cowan, 1984; Verbruggen & Logan, 2009; Verbruggen et al., 2019). In such tasks, a specific movement is initially instructed by a Go-signal. On some trials, salient stop-signals then instruct the subject to inhibit and cancel that movement prior to its execution. Stop-signals trigger activity within the right inferior frontal cortex (Swann et al., 2009), which in turn rapidly activates the STN (Chen et al., 2020). STN activity is then followed by thalamic signaling (Diesburg et al., 2021) and ultimately results in the inhibition of the motor cortex (Coxon et al., 2006; Wessel et al., 2022). Recent work has shown that inhibitory commands along this chain are sent through transient, burst-like signals in the β-frequency band (13-30Hz, Little et al., 2019; Wessel, 2020; Enz et al., 2021; Schaum et al., 2021; Choo et al., 2022).

Stop-signal tasks have contributed greatly to our basic understanding of response inhibition. However, they are a poor model for response inhibition in the real-world (Hannah & Aron, 2021). Real-world demands on the response inhibition system are highly specific and require a large degree of selectivity – both with respect to the stimuli that trigger inhibition (Bissett & Logan, 2014), and with respect to the movements that have to be inhibited (Aron & Verbruggen, 2008; Wadsley et al., 2022). First, unlike classic stop-signal tasks (which include a single unequivocal stop-signal) only specific salient stimuli in the real-world require response inhibition, whereas other do not. For example, pedestrians need to inhibit their walking movements after a car horn, but not after a ship’s air horn. Second, real-world inhibition is also ‘response-selective’. Rather than inhibiting all initiated movements (as in the classic stop-signal task), only some ongoing movements have to be stopped, whereas others have to continue. For example, pedestrians need to inhibit their walking movements without dropping their bag when hearing a car horn.

The study of this more realistic, stimulus- and response-selective response inhibition requires more complicated laboratory paradigms than the classic stop-signal task. Emerging work using such paradigms has shown that both stimulus- and response-selective stopping substantially alter the effects of response inhibition. Selective response inhibition changes both the behavioral stopping process (Aron & Verbruggen, 2008; Bissett & Logan, 2014; Bissett et al., 2021), as well as the physiological effects of response inhibition on the motor cortex (Duque et al., 2017; Tatz et al., 2021; Wadsley et al., 2023). This raises the question motivating the current study: how is this type of flexible, realistic response inhibition implemented in the fronto-subthalamic circuit?

Answering this question has been challenging, because complex, selective stopping tasks are difficult to implement in model organisms that allow precise measurements of subcortical activity (such as the STN). The subcortical nuclei of the basal ganglia that are paramount to response inhibition are small and embedded deeply within the brain (Jahanshahi et al., 2015; Forstmann et al., 2017). Measuring the activity of nuclei such as the STN with sufficient spatiotemporal precision requires invasive intracranial depth electrode recordings (Kühn et al., 2004). In the past, such recordings could only be performed in non-human animals (Fife et al., 2017; Schmidt et al., 2013) or in humans undergoing acute neurosurgery (Ray et al., 2012; Mosher et al., 2021). For these populations, it is difficult to perform complex, selective response inhibition tasks that mimic the requirements of real-world human environments. Rodents, for example, can only perform very simple versions of the classic, non-selective stop-signal task (Loyant et al., 2025). Humans can readily perform stimulus- and response-selective stopping tasks. However, intracranial recordings from the human STN (and other subcortical nuclei of the response inhibition network) typically have to be performed during acute neurosurgery. This is usually done in the context of deep-brain stimulation (DBS) surgery, which is a common treatment for several neurological and psychiatric disorders (Benabid et al., 1989; Hollunder et al., 2024; Lozano et al., 2008; Perlmutter & Mink, 2006). During DBS surgery, local field potentials can be recorded from the human STN (Marsden et al., 2001; Kühn et al., 2004). However, cognition and motor abilities are often substantially impaired during the perisurgical period and task performance windows are very short. So far, this has limited the work on response inhibition to non-selective stop-signal tasks.

To address this shortcoming, we here capitalize on the recent introduction of remote-sensing capable deep-brain stimulators (Cummins et al., 2021). These novel devices allow the wireless streaming of LFPs from the chronically implanted depth electrodes at any time after the surgery. With additional hardware and software (Soh et al., 2025), the chronic recordings from these DBS devices can be precisely aligned with behavioral tasks, as well as other neural recordings, with millisecond precision. This allows the measurement of LFPs from the human STN while the patients perform complex cognitive tasks, outside of the operating room, on their typical medication, in their optimal cognitive state, for extended periods of time (Soh et al., 2024; Hoy et al., 2024). Moreover, the extraoperative setting enables the collection of simultaneous cortical recordings (such as EEG or MEG) concomitantly with the subcortical LFPs. This makes it possible to study large-scale cortico-subcortical dynamics (Winkler et al., 2025).

Here, we performed a stimulus- and response-selective response inhibition task in 15 patients with DBS implants in the STN, while also recording EEG and EMG. This allowed us, for the first time, to our knowledge, to compare the cortico-subthalamic dynamics of selective and non-selective stopping in any species.

## Results

### Subjects and Behavior

Fifteen Parkinson’s Disease patients implanted with Medtronic Percept^TM^ sensing capable STN deep-brain stimulators performed a stimulus- and response-selective stopping task (**Figure 1A**) while undergoing a 64-channel scalp EEG recording, as well as EMG recordings from both responding muscles. Each trial began with a Go-stimulus that instructed the performance of an action. On 90% of trials, this action was a bimanual response, performed via button presses using the index fingers on both hands. On 10% of trials, a unimanual response was cued instead. These unimanual Go-trials were introduced as a comparison for the Selective-Stop trials, which required the continuation of a unimanual response. (In the following, we will refer to the regular, bimanual Go-trials as “Go-trials” and these rarer, unimanual Go-trials as “unimanual Go-trials”). Out of the 90% bimanual Go-trials, 30% featured a second, salient signal, with an adaptive delay. These salient visual stimuli either instructed the stopping of both responses (10% of trials), the stopping of one of the responses while the other continued (10%), or no stopping at all (10%). In keeping with the existing literature, we refer to the first type of trial – in which all movement has to be stopped – as “Global-Stop”-trials. We will refer to the second type of trial – in which one movement has to be stopped while the other has to continue – as “Selective-Stop”-trials. Finally, we will refer to the third type of trial – which features the same type of salient signal as the stop trials but does not require the stopping of any movement – as “Ignore” trials. These trials ensure that response inhibition is stimulus-selective.

**Figure 1.**
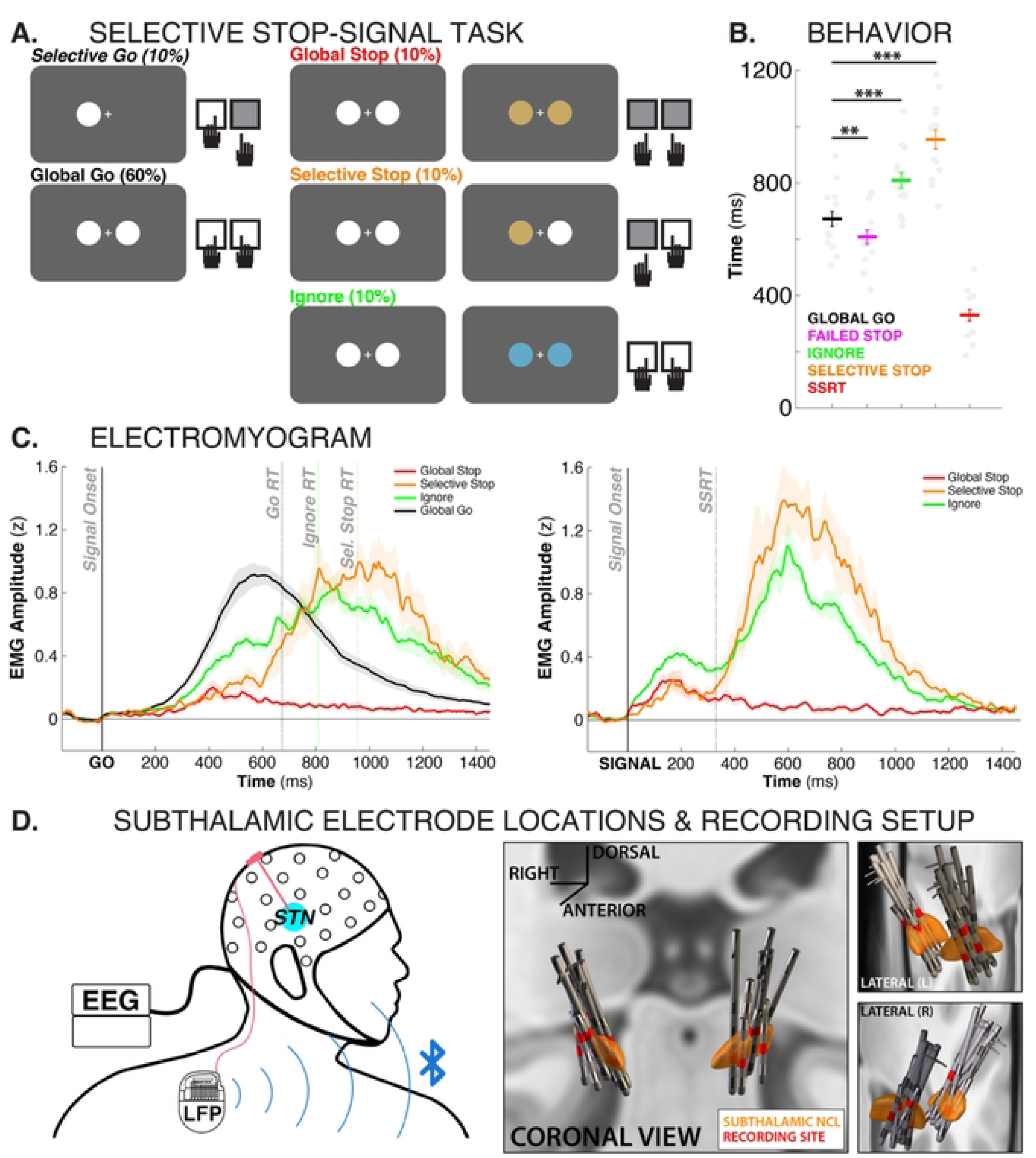
A) Stimulus- and response-selective stop-signal task diagram. B) Response times for the different conditions as well as SSRT. ** = p < .005, *** = p < .001. C) EMG traces time-locked to the Go-stimulus (left) as well as the different salient signals (right). D) Recording setup and 3D reconstructions of subthalamic depth electrode locations.

Subjects showed behavior typical of older adults (**Figure 1B**). Go-trial reaction time was 673ms (SEM: 27ms) on bimanual Go-trials and 711ms (SEM: 35ms) on unimanual Go-trials, with no significant differences between the two. All subjects showed good stopping performance, with an average stop-signal reaction time (SSRT) of 330ms (SEM: 20.1). As expected for the stop-signal task, failed stop trials (609ms, SEM: 25ms) showed consistently faster responses than Go-trials (t(14) = 3.4, p = 0. 0048, d = 0.6). Compared to bimanual Go-trials, both Selective-Stop trials (955ms, SEM: 34ms) and Ignore trials (809ms, SEM: 28ms) showed a significant slowing of reaction times of the continuing response (t(14) = 16.36, p <.0001, d = 2.2; t(14) = 9.08, p <.0001, d = 1.2). EMG traces for all trial types can be found in **Figure 1C**.

LFP data from the subthalamic depth electrodes (**Figure 1E**) were downloaded from the implanted DBS device and precisely aligned with the EEG recording using established methods (Soh et al., 2025). The digital event information from the EEG recording was then used to inform the analysis of the LFP trace, as was done in previous studies (Hoy et al., 2024; Soh et al., 2025).

### Cortical activity shows early overlap and later divergence depending on selectivity

We trained several multi-variate decoders to distinguish whole-scalp EEG patterns associated with the different critical trial types. Activity after salient signals (Global-Stop, Selective-Stop, Ignore) was decodable from activity after Go-signals very early after the respective signal (see red, orange, and green lines in **Figure 2A**, *p < .01, cluster-corrected for multiple comparisons*). However, differences between the three salient signal types themselves could not be detected until substantially later in the trial. Starting at ∼450ms, all three trial types showed consistent and decodable differences from one another. The topographical voltage map during this later time period (specifically, the 600 to 666ms period during which all three processes showed significant pairwise decoding differences) showed a centrally-located voltage positivity that was stronger on Global-Stop compared to both Ignore and Selective-Stop trials, and was also stronger on Selective-Stop compared to Ignore trials. This activity represents the classic stop-signal P3 event-related component (de Jong et al., 1990; Hervault et al., 2025), which typically peaks at electrode Cz. The ERP at that electrode is therefore also shown in **Figure 2A**.

**Figure 2.**
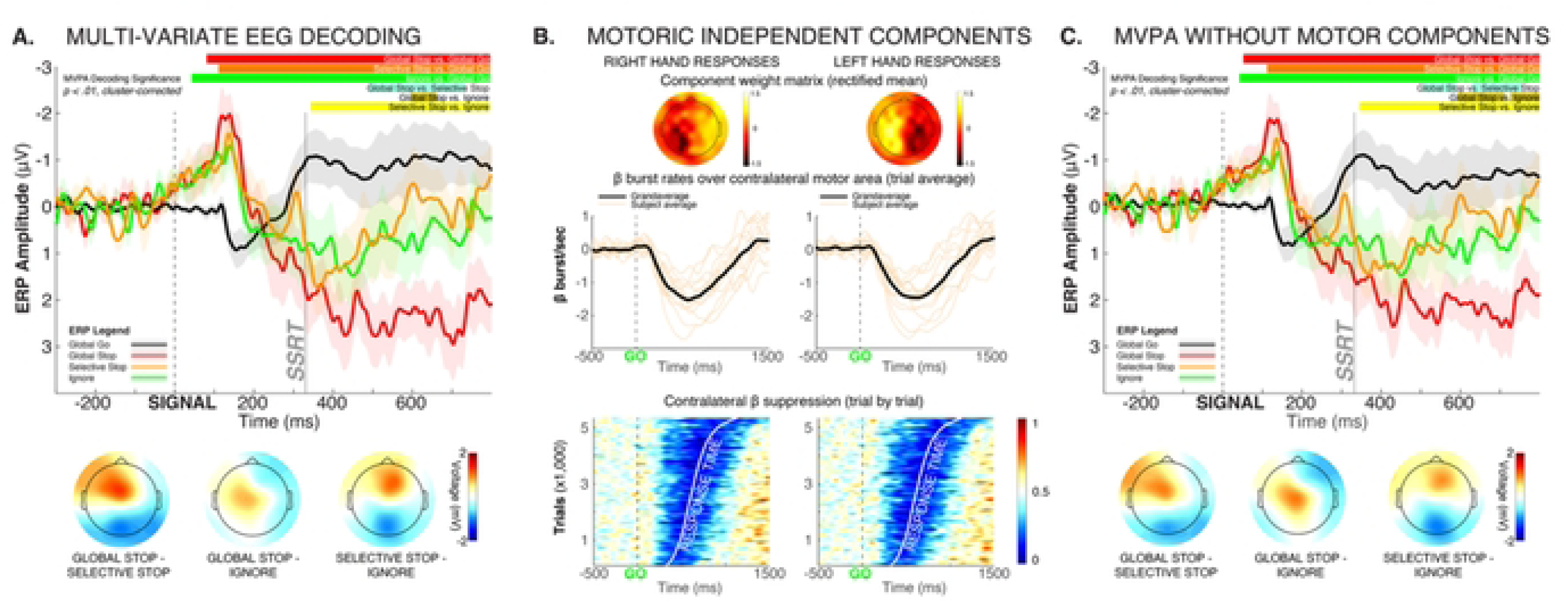
Multi-variate decoding of whole-scalp EEG data shows early processing overlap between Global-Stop, Selective-Stop, and Ignore trials, with only later differences emerging. A) Significant periods of multi-variate decoding for the different contrasts are shown on top of the plot (p < .01, cluster-corrected for multiple comparisons). The voltage topographies in the time period from 600 to 666ms are shown at the bottom, as this time period showed significant pairwise differences in decoding between all three conditions with salient signals. Since these topographies show that the EEG differences center around electrode Cz, the event-related potential at that electrode is depicted in the line graph above. B) To ensure that these late differences cannot be accounted for by motor activity, we identified the independent EEG components in each subject that contained response-related motor activity. The topographical plots show the mean rectified component weight matrix for right- and left-hand motor responses, respectively. Hotspots over contralateral motor areas can be clearly seen. Moreover, each subjects’ selected component showed reductions of β bursting after the onset of the Go-signal (orange lines in middle plots), consistent with established EEG motor signatures. Finally, the contralateral β activity suppression reflected in the selected components clearly tracked response times, as revealed by the ERP Image at the bottom of Figure B. C) Same analyses and plots as A, but after removal of the components shown in B from the compound EEG signal mixture.

We then ensured that this later difference was not merely a consequence of the different types of motoric activity between the three conditions (**Figure 2B-C**). After removing motoric signal components from the compound EEG signal using independent component analysis (ICA, **Figure 2B**), we retrained the same multi-variate decoders as in **Figure 2A**, with very similar latencies emerging from the analysis (**Figure 2C**, *p < .01, cluster-corrected for multiple comparisons*).

Together, these EEG decoding analyses suggest that at the cortical level, processing between Global-Stop, Selective-Stop, and Ignore trials is initially indistinguishable. Significant differences in processing emerge later in the trial, chiefly as differences in processing over fronto-central electrode sites.

### STN activity also shows early overlap and later divergence depending on selectivity

The full-spectrum STN LFP condition comparisons can be found in **Figure 3A**. In line with prior work, inhibitory STN activity largely took place in the β frequency band (Kühn et al., 2004; Ray et al., 2012; Diesburg et al., 2021). β-band activity in the full-spectrum analysis was increased, alongside low-frequency activity, on Global-Stop compared to Go trials (**Figure 3A**, left panels, p < .05, *cluster-corrected for multiple comparisons*). STN β-band activity in particular was also increased on Global-Stop compared to Selective-Stop trials, as well as compared to Ignore trials (**Figure 3A**, middle and right panels). As prior work has found, STN β-band activity takes the form of discrete, transient ‘burst-like’ events (Diesburg et al., 2021; **Figure 3C**). **Figure 3B** shows the β burst probability timelines within the bilateral STN for each trial type, as well as the baseline burst rate in the 500ms period prior to the Go-signal (dashed line). At the time of the signal, each trial type showed a declining rate of β bursts relative to baseline. This reflects the disinhibition of the motor system during the initiation of the movement. (Note that the Go trial data was time-locked to a time point that matched the current stop-signal delay, rather than the Go-signal itself, to account for this activity). Shortly after the presentation of the salient signal, however, all three trial types (including Ignore trials, in which the signal did not instruct any stopping) showed a fast-latency increase in STN β burst rates relative to go-trials. STN burst rates returned to pre-trial baseline prior to SSRT in all three trial types that featured salient events (Global-Stop, Selective Stop, and Ignore). This fast-latency increase in STN activity, which occurred regardless of stopping requirements, is in line with the STN’s role in implementing rapid, short-lasting and broad motor inhibition after any type of salient event (Wessel & Aron, 2017).

**Figure 3.**
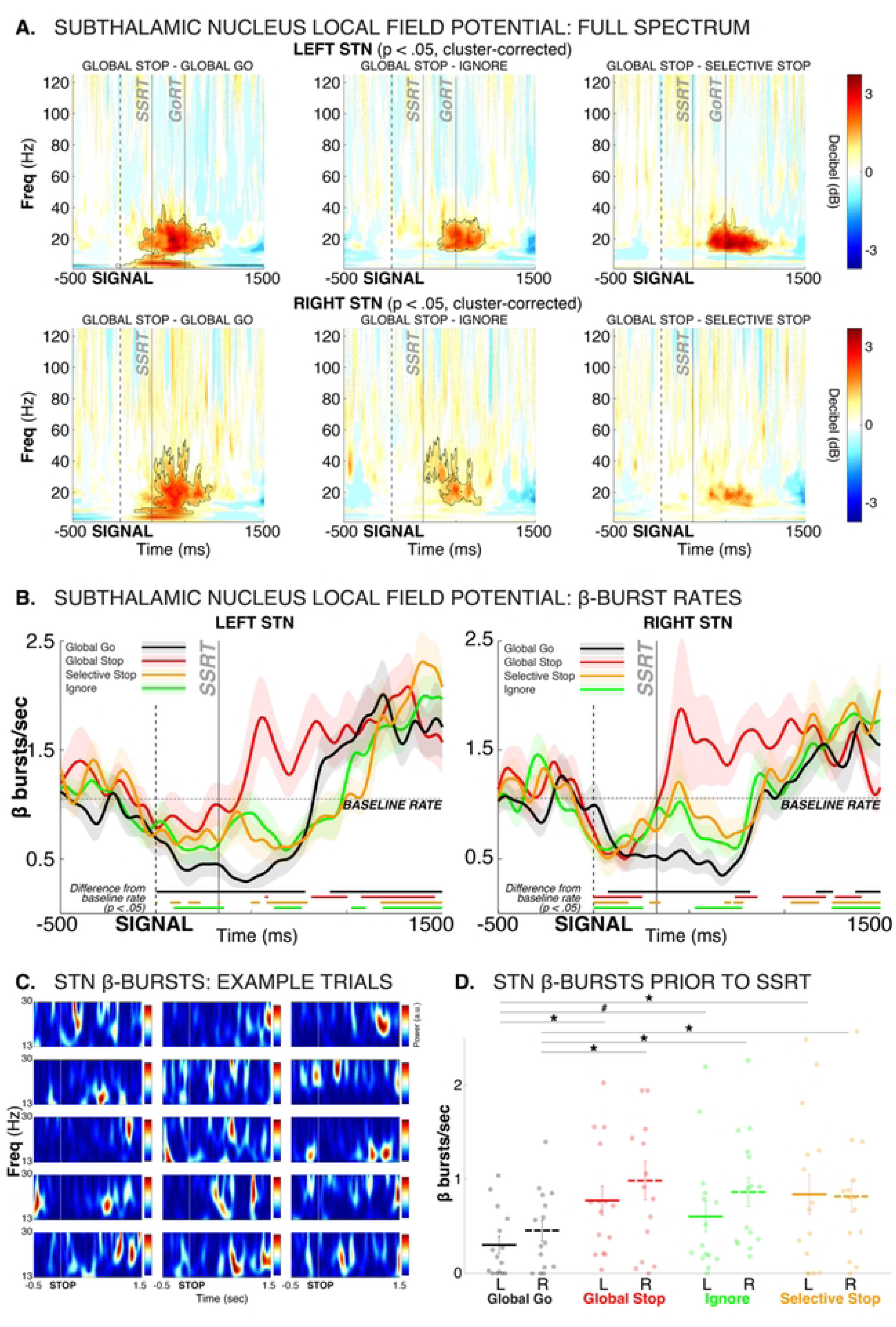
Subthalamic nucleus dynamics during selective and non-selective stopping. A) Full-spectrum event-related perturbation conditional comparisons between Global-Stop and Go trials (left), Global-Stop and Ignore trials (middle) and Global-Stop and (Response-)Selective-Stop trials. Black outlines denote statistical significance at p < .05, cluster-corrected for multiple comparisons. Separately for left and right STN. B) β burst probability time series, split by condition, separately for left and right STN. Solid lines at the bottom of each plot denotes statistical differences of the same-colored curve from baseline (p<.05). Go trial burst timeline is time-locked to the current stop-signal delay to account for differences in motor initiation processes between Go trials and Stop/Ignore trials. Baseline rate (dashed line) is the burst rate in the 500ms prior to the Go-signal in the respective STN. C) Individual Global-Stop trial examples from each subject, showing that STN β-band activity is contained in transient, discrete burst events, rather than as an amplitude-modulated continuous oscillation. D) β burst counts in the pre-SSRT period (50ms prior to SSRT to end of SSRT, individualized for each subject), per condition. * = p < .05, # = p < .05 (one-sided).

Prior computational modeling work and single-neuron recordings from non-human primates have suggested that motor-system inhibition during action-stopping begins just prior to the end of the SSRT period (Boucher et al., 2007). In line with this, STN β burst rates were increased relative to Go trials for all three trial types right before the end of SSRT (**Figure 3D**).

Most notably, substantial condition differences then emerged after SSRT. On Global-Stop trials, where all movement had to be stopped, STN β bursting exceeded the baseline burst rate for a sustained period of over a second (**Figure 3C**). In contrast, both the Selective-Stop condition (where only part of the response had to be inhibited) and the Ignore condition (where no inhibition was necessary at all) featured an immediate reduction of STN β bursting back to below-baseline levels. Notably, after the response was made, all three trial types with responses (Selective Stop, Ignore, and Go-trials) then showed a similar above-baseline increase in STN β burst rates to that observed immediately after SSRT on Global-Stop trials.

Together, this suggests that all salient signals initially trigger a fast implementation of response inhibition. When non-selective inhibition of all ongoing responses is required (global-Stop), the motor system is then inhibited broadly and for an extended period of time, signified by a sustained above-baseline β bursting in STN. In contrast, when only partial, response-selective response inhibition (or no inhibition at all) is required, this pattern is delayed until after the continuation of the appropriate response. Prior to that, the STN-based inhibition is re-released to enable the execution of the respective response on those trials. This *inhibit-release-inhibit* dynamic on Selective-Stop and Ignore trials also explains why there is substantial behavioral slowing on trials with Selective-Stop and Ignore trials (**Figure 1C**).

### β activity before and after SSRT is tightly coupled between frontal areas and STN

As highlighted above, both STN and cortical activity suggests that Global-Stop, Selective-Stop, and Ignore trials initially feature a shared process. Critical differences between the conditions were found only later in the trial (after SSRT). On the scalp, this was signified by unique activity over fronto-central areas (**Figure 2A, C**). In the STN, this was evident from sustained, above-baseline STN activity (**Figure 3B**). These scalp and STN signals shared a similar time course, suggesting a potential relationship between cortical and subthalamic processing.

Indeed, STN β band activity on trials with salient signals (Global-Stop, Selective-Stop, Ignore), was highly correlated with β activity over fronto-central scalp regions, both shortly before and immediately after SSRT (**Figure 4A**; *electrode sites and time windows with significant cross-area correlations are highlighted at a threshold of p < .05, corrected for multiple comparisons*). STN β band activity in the time period immediately prior to SSRT (the last 50ms before subject-specific SSRT, see **Figure 3D**) was predicted by β band activity at fronto-central electrodes in the time window starting around 100ms prior to SSRT. Similarly, STN β-band activity in the time window immediately after SSRT was correlated with fronto-central EEG in the post-SSRT period. **Figure 4B** shows the same relationship with better temporal resolution. EEG β-band activity in the fronto-central ROI was correlated with bilateral STN β-band activity for an extensive time period both before and after SSRT (**Figure 4B**; *black outline shows single-trial significant correlations at a cluster-corrected critical p-value of 0.0027 – i.e., .05 divided by 18, the number of spatial ROIs in*). Note that the ‘motor’ electrodes C3 and C4 were contained in the lateral ROIs, suggesting that this cortico-subthalamic correlation did not merely reflect the general β coupling between STN and motor cortex (see **Figure 5A** below).

**Figure 4.**
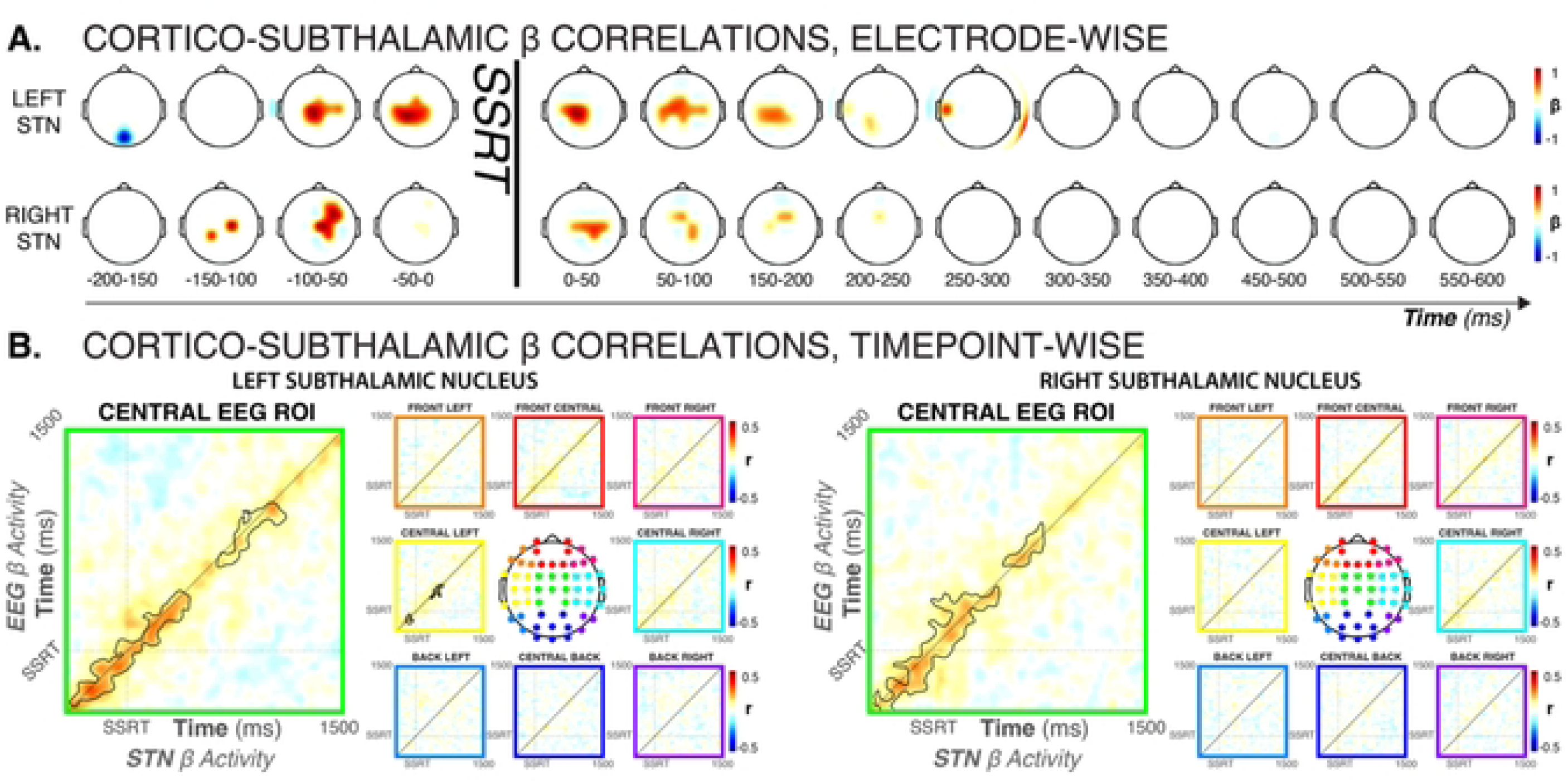
Cortico-subthalamic β-band dynamics across all trials with salient signals (Global-Stop, Selective-Stop, Ignore) showed strong interactions between bilateral STN and fronto-central brain regions. A) Single-trial β-band amplitude correlations between left and right STN activity before and after SSRT and whole-scalp, shown for each individual electrodes, thresholded for significance at p < .0063 for the pre-SSRT windows (i.e., .05 divided by 8, the number of time windows) and p < .0025 for the post-SSRT windows (i.e., .05 divided by 20, the number of time windows). Plots prior to SSRT show correlations between scalp activity in the indicated time window and STN activity in the 50ms window prior to SSRT. Plots after SSRT show correlations between scalp activity in the indicated time window and STN activity in the 100ms window after SSRT. B) Time-lagged trial-wise correlations between nine EEG ROIs and left/right STN β-band activity. The central ROI (electrodes FC1, FC2, FCz, C1, C2, Cz, CP1, CP2, Cpz) showed strong correlations with β-band activity (black outline denotes cluster-corrected significance at a cluster- and sample-wise p = = .00027) both before and after SSRT. No other ROI shows significant correlations for both left and right STN activity.

**Figure 5.**
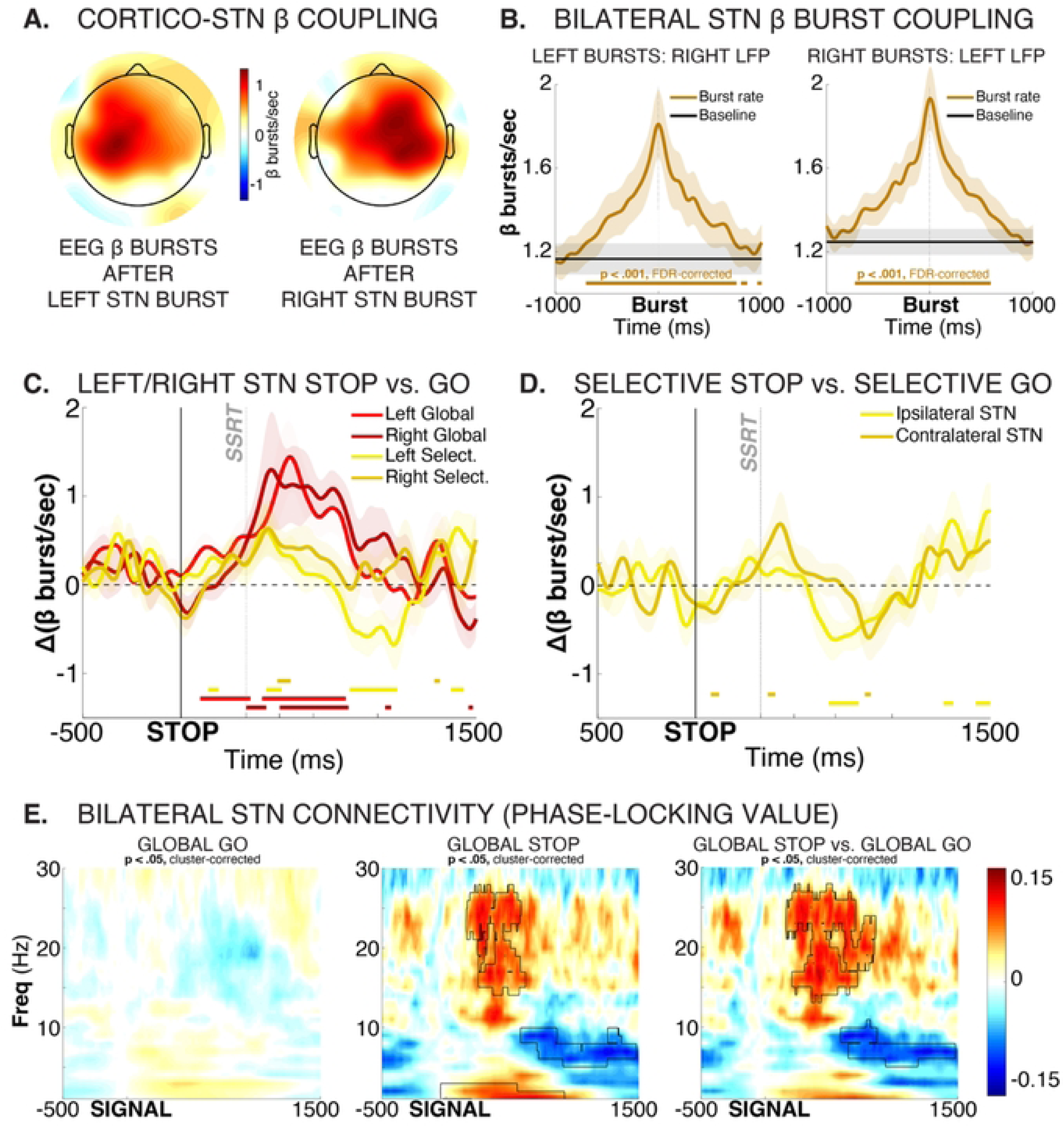
Left and right STN show highly coupled activity during response inhibition, with little evidence towards strong lateralization. A) Topographical representation of EEG β burst rates immediately (0 - 50ms) after STN bursts, separately for left and right STN. Plot is thresholded at p < .00081 (i.e., p < .05 divided by the number of channels to correct for multiple comparisons). A clear ispilateral organization of relative burst timing between STN and (motor) cortex is evident. B) β bursts are tightly coupled across both STNs, with right STN burst probability significantly increased before and after left STN bursts (left plot) and vice versa). Brown solid line at the bottom of each plot shows significant differences in burst rate from baseline burst rate (p < .001, FDR-corrected for multiple comparisons). C) Left and right STN show similar dynamics during Global and Selective response inhibition. Plot shows STN condition-difference β burst probability timelines between Global-Stop and Go trials, separately for left (bright red) and right (dark red) STN, as well as condition-difference β burst probability timelines between Selective-Stop and Go trials, separately for left (bright yellow) and right (dark yellow) STN. Solid lines on the bottom of the plot show significant differences from zero for each curve at each time point. No significant differences between the left and right STN condition comparisons survived FDR correction for multiple comparisons, neither for Global nor for Selective response inhibition. D) Contra- and ipsilateral STN show similar dynamics during Selective response inhibition. Plot shows STN condition-difference β burst probability timelines between Selective-Stop and unimanual Go trials, separately for left (bright yellow) and right (dark yellow) STN. Solid lines on the bottom of the plot show significant differences from zero for each curve at each time point. No significant differences between the contra- and ipsilateral STN condition comparisons survived FDR correction for multiple comparisons. F) During Global response inhibition, β-band connectivity (phase-locking value) between left and right increased significantly, suggesting that both STNs coordinate the complete, non-selective cancellation of all initiated movements. Black outline denotes significant differences at p < .05 (cluster-corrected for multiple comparisons).

These analyses show that the β band dynamics between fronto-central EEG signals and bilateral STN were highly correlated during the specific adjustments to behavior necessitated by the salient signal (i.e., global inhibition, selective inhibition, or continuation).

### No evidence of lateralized STN activity during response inhibition

Finally, whether response inhibition activity in STN is lateralized has been a contentious question. Notably, an influential fMRI study reported that STN activity during (Global) response inhibition is lateralized to the right STN, regardless of the side of the action that has to be stopped (Aron & Poldrack, 2006). However, anatomical connections (Coudé et al., 2018) and functional coupling between STN and motor cortex typically show strong ipsilateral organization (Litvak et al., 2011; Werner et al., 2025). Consequently, unilateral STN lesions typically lead to clinical symptoms affecting contralateral limbs, such as hemiballismus (Posturna & Lang, 2003). Indeed, DBS electrodes are sometimes implanted only contralateral to the most affected side (Kumar et al., 1999). Thus, one could predict that each STN should specifically inhibit responses made with the contralateral hand. For that reason, some intracranial STN recording studies of the classic stop-signal task have only recorded from the contralateral STN (Ray et al., 2012; Mosher et al., 2021). Others have averaged data from both STN (Bastin et al., 2014; Benis et al., 2014; Diesburg et al., 2021). However, no systematic investigations of STN LFP lateralization during response inhibition exists. Our data allow such an investigation.

First, in line with previous work, cortico-subthalamic dynamics throughout the recording were strongly lateralized to the same side. In the EEG, increases in β burst rate that took place after STN bursts clustered tightly around ipsilateral motor sites (**Figure 5A**). Very notably, β dynamics between the two subthalamic nuclei were also highly correlated throughout the recording. β burst rates in left STN were strongly increased both before and after individual bursts in right STN, and vice versa (**Figure 5B**).

With regards to response inhibition, both left and right STN showed comparable increases during both selective and global response inhibition (**Figure 5C**). Left, rather than right STN showed qualitatively earlier effects between Global/Selective-Stop trials and Go trials (see also full-spectrum plots in **Figure 3A**). However, this difference did not reach statistical significance at any time point for either contrast (selective/non-selective response inhibition). Either way, there was little evidence towards a right-lateralization of response inhibition at the level of the STN.

During selective stopping, the difference between Selective-Stop and unimanual Go trials was slightly more pronounced for the STN contralateral to the movement, with the contralateral STN showing significant early condition differences between Stop and Go trials that were absent in ipsilateral STN (**Figure 5D**). This is in line with the ipsilateral dominance in coupling between STN and motor cortex (**Figure 5A**). However, the condition difference between contralateral and ipsilateral contrasts was not significant above FDR-correction at any point.

In sum, there were no strongly lateralized effects for either type of response inhibition in the STN. Moreover, STN β activity was tightly coupled throughout the recording, which suggests that both nuclei may be working in tight coordination. Indeed, during Global-Stop trials, significant phase-locking emerged between both STN, suggesting that both nuclei work together to effect non-selective response inhibition (**Figure 5C**).

## Discussion

To our knowledge, this is the first investigation of the cortico-subcortical dynamics of selective and non-selective response inhibition. We used a stimulus- and response-selective response inhibition task that required participants to either inhibit all ongoing responses (a ‘Global’ stop condition that matches the non-selective inhibitory requirements of the classic stop-signal task), inhibit one of multiple responses while continuing with another (response-selective stopping), or to ignore a similar salient signal and continue all responses (to ensure that response inhibition is stimulus-selective).

All three salient stimuli rapidly led to activity changes in the cortex and the subthalamic nucleus. Notably, neither the initial subthalamic nor the initial cortical activity differed between the three critical conditions. EEG decoding did not reveal any unique early activity that distinguished the three types of trials (**Figure 2A,C**), and STN β-frequency burst rates increased back to baseline levels rapidly after all three types of signals (**Figure 3B**). In line with this, Ignore and Selective-Stop trials – both of which required the continuation of one or both movements – featured a notable reduction in the EMG of the responding effector at the same time as the Global trials (**Figure 1C**). Together, these findings show that – regardless of inhibitory requirements – all salient events trigger a rapid and transient re-inhibition of the motor system. This aligns with past findings of broad suppression of the excitability of the cortico-spinal tract after any type of salient signal, regardless of whether they require global stopping (Coxon et al., 2006; Badry et al., 2009), selective stopping (Wadsley et al., 2023), or no stopping at all (Wessel & Aron, 2017).

After this early inhibitory process, substantial differences between the conditions emerged. On Global-Stop trials, which required the non-selective inhibition of all active responses, the STN showed a sustained, above-baseline β bursting rate that lasted for at least more than one second past the initial return to baseline (**Figure 3B**). In comparison, on Selective-Stop trials and Ignore trials, STN β bursting reduced below baseline again during this time period. This likely reflects a release of inhibition to allow for the execution of the continued response.

On the cortical level, unique patterns of fronto-centrally distributed scalp activity emerged during this same time period, as revealed by multi-variate EEG decoding. Single-trial correlation analyses across the three critical conditions showed a tight relationship between this fronto-central cortical activity and subthalamic activity throughout this period (**Figure 4**). While scalp EEG does not allow for a precise localization of this signal to a specific part of the cortex, we speculate that this cortico-subcortical relationship reflects an interaction between the STN and the cortical pre-supplementary motor area (preSMA). The preSMA is a key node in the response inhibition circuit (Nachev et al., 2007; Roberts & Husain, 2015; Wolpe et al., 2022; Apšvalka et al., 2022). The preSMA and STN are anatomically connected (Inase et al., 1999; Hollunder et al., 2024; Oswal et al., 2021), and the integrity of this connection is directly related to the efficacy of response inhibition (Forstmann et al., 2012; Coxon et al., 2012). Finally, intracranial (Swann et al., 2012) and MEG studies (Schaum et al., 2021) have shown that preSMA activity during response inhibition takes place in the β frequency band. As such, our finding that fronto-central cortical activity is tightly related to subthalamic β activity during trials with different response inhibition requirements is consistent with the idea that an inhibitory preSMA-subthalamic pathway governs the precise implementation of context-specific response inhibition.

In addition, our unique, bilateral intracranial STN recordings allowed us to test the purported lateralization of STN activity during response inhibition (Aron & Poldrack, 2006). First, we confirmed that coupling between STN and motor cortex was highly lateralized (see also Litvak et al., 2011; Werner et al., 2025), as STN β bursts were followed by significantly increased bursts rates over ipsilateral motor cortex. Second, we found that across the entire recording, burst timing was highly concurrent across both STN. Third, we found no evidence for the purported right lateralization of STN activity during either type of response inhibition (Selective or Global). Neither did we find strong evidence to support a highly specialized role for the contralateral STN in unimanual (selective) response inhibition. Indeed, during non-selective response inhibition, strongly coherent β band activity emerged between both STN. This suggests that response inhibition is carried out by both STN, perhaps in tight bilateral coordination, though future work should systematically assess the role of handedness and response laterality in these effects.

Together, this study shows that a cortico-subthalamic circuit consisting of fronto-central cortical areas and the bilateral subthalamic nuclei exerts flexible inhibitory control over behavior. Selective inhibition is achieved via a two-step dynamic. First, any salient signal that occurs during response preparation automatically instantiates a short-latency baseline-return of STN β burst rates. This countermands the initial reduction of STN β bursting that signifies the release of inhibition during responding. Second, a coordinated and flexible fronto-subthalamic process then implements specific adjustments to the inhibitory activity of the bilateral STN. The resulting STN activity is highly contingent on the control requirements of the specific situation. When responses have to continue (e.g., during selective stopping), inhibition is released once more, signified by a return of STN β bursting below baseline. When all responses have to be inhibited (i.e., during Global-Stopping), the STN immediately enters a state of sustained, above-baseline β bursting, likely in an effort to quell any residual activation of the response (see Schmidt & Berke, 2017; Diesburg & Wessel, 2021 for recent theoretical models of these dynamics).

This sustained, above-baseline STN activity during non-selective response inhibition has substantial implications for the basic science of response inhibition. The influential horse-race model of action-stopping (Logan et al., 1984) suggests that stop-signal reaction time (SSRT) denotes the end of the ‘race’ between the prokinetic Go process (response execution) and the antikinetic Stop process (response inhibition). Decades of subsequent work on the neuroscientific basis of response inhibition has therefore assumed that neural processes critical to response inhibition have to occur before the end of SSRT. Recent work has challenged this notion, showing that substantial amount of inhibitory activity continues to take place after SSRT (e.g., de Navas et al., 2025). Indeed, in the current study, the initial inhibitory activity that took place prior to SSRT did not return EMG activity back to baseline (**Figure 1B**, see also Tatz et al., 2021 for similar findings in healthy young adults). Instead, EMG following both Ignore and Selective-Stop signals remained significantly above baseline after the initial reduction. Even Global-Stop-trials, on which no overt response is made, featured substantial above-baseline EMG activity well after the initial suppression of the EMG and SSRT (**Figure 1B**). Together with the sustained, above-baseline STN activity this suggests that additional inhibitory activity persists for an extended period after SSRT if responses are to be fully inhibited. We speculate that this is because – unlike assumed in the horse-race model – it is not sufficient for response inhibition to initially ‘win the race’ against response execution. Instead, it has to ‘keep winning’ for an extended period of time to prevent a belated execution of the inappropriate response (Schmidt & Berke, 2017; Diesburg & Wessel, 2021). Future work is needed to test this hypothesis directly.

In sum, we found that a cortico-subthalamic circuit exerts flexible response inhibition depending on the contextual requirements on inhibitory control – similar to those found in the real world. Once a salient signal occurs, all responses are partially inhibited. During selective stopping, this broad inhibition is subsequently released to enable the continuation of the appropriate response. In contrast, during non-selective stopping, the motor system is over-inhibited for an extended period of time, signified by a sustained, above-baseline rate of burst-like β activity in the STN.

## Materials and Methods

### Participants

All experimentation was performed in accordance with the Declaration of Helsinki. All experiments were clinically supervised by an attending neurologist with subspecialty training in Parkinson’s disease (EYU or QZ). Participants signed a consent form approved by the University of Iowa IRB board (IRB ID: 202002751). Enrollment in the study was contingent upon scoring greater than 19 on the Montreal Cognitive Assessment (MoCA) and screening by an attending neurologist with subspecialty training in Parkinson’s disease (EYU). We recruited sixteen Parkinson’s Disease patients who had been previously implanted with wireless sensing capable Percept PC DBS stimulators (*Medtronic plc., Minneapolis, MN, USA*) targeting the bilateral STN (Mean Age: 67.5y; Age Range: 53-79; 4 female; 2 left-handed; for clinical characteristics see **Supplementary Table 1**). Participants were recruited from JDWG’s neurosurgical practice. Two patients had stopped taking dopaminergic Parkinson’s Disease medications due to the effectiveness of the DBS treatment. The remainder of the patients performed the task on their routine medications to allow for ideal cognitive and motor performance. One participant did not complete the experiment due to an inability to perform the experimental task.

### Experimental task

The task comprised 12 blocks that were completed within a single experimental session for each patient. Of the 12 blocks, two included 42 Go trials each, and the remaining ten contained 60 trials each, comprising both Go and signal trials (684 trials in total). On Go trials (70%), the patients were instructed to rapidly execute a response to a Go stimulus, whereas on signal trials (30%), a salient signal could appear after the Go stimulus, instructing the patients to either continue the response, stop the full response, or stop part of the response (**Figure 1A**).

On Selective-Go trials (10%), a white circle appeared on the left or right side of the central fixation cross, prompting the patients to quickly perform the corresponding left- or right-index-finger button press. On Global-Go trials (60%), two white circles appeared on the left and right sides of the fixation cross, prompting patients to perform a bimanual response with both index fingers.

On Global-Stop trials (10%), the Global-Go stimulus was followed by a Global-Stop signal, consisting of both white circles turning gold. In this case, the patients were instructed to abort the bimanual response and withhold any button press. In Selective-Stop trials (10%), the global-Go stimulus was followed by a Selective-Stop signal, consisting of one of the two circles (either left or right) turning gold while the other remained white. In this case, the patients were instructed to abort the response on the signaled side while continuing the response on the other side. On Ignore trials (10%), the Global-Go stimulus was followed by an Ignore signal, consisting of both white circles turning blue. In this case, the patients had to continue the bimanual response and ignore the salient signal. In Global-Stop and Selective-Stop trials, the stop-signal delay (SSD; i.e., the interval between Go and Stop signals) was initially set to 200ms and was then dynamically adjusted according to each patient’s stopping performance. This procedure was implemented separately for Global-Stop, left Selective-Stop, and right Selective-Stop signals: the SSD increased by 50ms after successful stopping (no response) and decreased by 50ms after failed stopping (response performed despite the stopping instruction). This tracking procedure was intended to converge on approximately 50% successful stopping for each stop type (Verbruggen & Logan, 2009; Verbruggen et al., 2019). Ignore trials used the current value of the Global-Stop SSD but did not modify it.

Each trial began with a central fixation cross presented for 1000ms. The Go stimulus remained visible for up to 1,000ms, and responses were accepted within a 1,500ms deadline from Go onset. The total trial duration was 4,000ms. Before the main task, the patients performed a practice block of 60 trials, which was repeated if their overall stopping success rate fell outside the 0.25–0.75 range. The assignment of gold versus blue colors to the Stop versus Ignore conditions was randomized across participants.

The experiment was programmed using Psychtoolbox 3 on MATLAB 2017a, running under Ubuntu Linux. Stimuli were displayed on a BenQ XL2420B monitor featuring a 120 Hz refresh rate and a resolution of 1920×1080 pixels. Manual responses were captured using a standard QWERTY USB keyboard, except in two cases, in which a USB response device (Kinesis Savant Elite 2) was used.

### Behavioral Analysis

We estimated SSRT using the integration method with replacement of Go omissions (Verbruggen et al., 2019). SSRT indexes stopping speed: a longer SSRT indicates slower stopping, and a shorter SSRT indicates faster stopping (Logan & Cowan, 1984). It was computed using the probability of responding in global stop-signal trials and the global Go RT distribution. Prior to SSRT calculation, we checked whether the subjects’ behavior was suitable for computing SSRT. First, we verified the efficacy of the staircasing procedure used in the task (mean stop probability: 53%; range: 47-60%). Second, we tested whether the Go RT was slower than the failed-stop trial RT to confirm that the prediction of the independent horse-race model was met, using a paired-samples t-test.

We then quantified the average RT for each condition. To test whether signal trials yielded longer RT than Go trials, we compared global Go RT with selective stop-signal RT and global Go RT with ignore-signal RT using paired-samples t-tests. All behavioral analyses used trials from blocks in which both Go and signal trials were intermixed. Trials with RTs under 250ms were discarded.

### LFP, EEG, and EMG recording

The Percept PC Neurostimulator’s communicator device (Medtronic model 8880T2) was used to wirelessly stream STN LFP data to the provided programmer tablet (Model CT900, *Samsung Galaxy Tab S5e*) via Bluetooth at 250 Hz (the maximum available frequency). The two directional sensing DBS leads are equipped with four rows of depth electrodes each, spaced 1.5 mm apart. LFP recordings were made from bipolar contact montages above and below the clinical stimulation electrode(s) on each side, thus centering the recorded voltage on the clinically efficacious contact site. The data were online filtered using a lowpass filter of 100 Hz and a highpass filter at 1 Hz. To ensure a stable wireless connection, both the communicator and the operating tablet were consistently kept within a 50 cm range. Participants performed the experiment with the DBS stimulator in the off state to enable the wireless sensing feature and suspend stimulation of the STN. An attending neurologist verified the DBS parameters before and after the experiment to ensure that participants left with their original DBS configurations and their DBS enabled.

EEG was acquired using a 64-channel EEG cap, amplified via Brain Vision MRplus amplifiers (*Brain Products, Garching, Germany*). The sampling rate was set to 5kHz to enable a precise alignment between EEG and LFP data and to ensure sufficient sampling rate for EMG data acquisition. The EEG signal was online filtered at a low cutoff .01 Hz and a high cutoff of 1000 Hz. The reference electrode was Pz, and the ground was Fz. EMG was recorded through the EEG amplifier from each hand using two electrodes placed on the belly and tendon of each FDI muscle. Ground electrodes were placed on the distal end of each hand’s ulna.

Before each block of the task was started, the DBS was activated for ∼12 seconds and then turned off again. The next block did not start until 12 seconds after the DBS was turned off. This procedure introduced simultaneous artifacts in both the EEG and LFP traces, which were used to precisely align the EEG and LFP datasets (Soh et al., 2025). After successful alignment, the LFP data were analyzed using the digital event information from the EEG system.

### STN electrode contact reconstruction

DBS electrode location was reconstructed from a pre-operative anatomical MRI scan and a post-operative MRI scan (1 patient) or computed tomography (CT) scan (all other patients), both of which are conducted as part of the standard clinical implantation procedure. Reconstructions were performed via LeadDBS using the standard steps and settings (Horn & Kühn, 2015). First, linear co-registration of the post-operative scans to the pre-operative T1-weighted images was performed via ANTs and then refined to account for potential intra-operative brain shift through the LeadDBS subcortical refinement module. Normalization warp fields were then refined manually using LeadDBS’ WarpDrive tool. For CTs, LeadDBS’ PaCER implementation was then used for the final reconstruction. For MRIs, LeadDBS’ TRAC/CORE algorithm was used. Two patients’ electrode locations could not be reconstructed due to the lack of high-resolution post-op imaging.

### EEG/LFP/EMG preprocessing

All electrophysiological data were processed using custom scripts coded with MATLAB 2023b (The MathWorks), which primarily relied on the features of the EEGLAB toolbox (Delorme & Makeig, 2004). After alignment, all data were resampled to 500Hz (EEG/EMG: from 5k to 500 Hz; LFP: from 250 to 500Hz) and bandpass filtered (EEG: 0.5-49Hz; EMG: 10-40Hz; LFP: 0.5-100Hz). The continuous EEG data were visually inspected to identify segments with non-stereotypical artifacts (e.g., muscle artifacts, line noise, etc.) that were removed from all EEG, EMG, and LFP data. The EEG data were re-referenced to the common average and then subjected to the independent component analysis (ICA; Makeig et al., 1996) to identify and remove stereotypical artifacts (blinks, saccades, and electrode artifacts).

### EMG analysis

Epochs were extracted from −200 to 1500ms relative to the events of interest. The EMG data were converted to RMS power by sliding a time window of ±5ms throughout the data, and then baseline-corrected by dividing the entire epoch by the mean of a 200-ms pre-stimulus window. The resultant data were then standardized across all conditions and time points using z-transformation for each hand separately. For Global trials, EMG recordings were averaged across hands. For Selective-Stop-signal trials, we focused on the continuing hand (e.g., left-hand EMG for right-hand stopping trials).

Global-Stop-signal EMG data included only trials with partial response EMG (prEMG; Raud et al., 2022). The individual prEMG threshold was determined by averaging a 200-ms window following stop-signal onset across trials for each subject. The maximum peak amplitude of this trial-averaged data was taken as the individual prEMG threshold. Any trials with EMP peaks that exceeded this threshold were included in the stop-signal trial EMG analysis.

### Multi-variate EEG decoding analysis

For the EEG decoding analysis, we used the ADAM toolbox (Fahrenfort et al., 2018). Whole scalp EEG data were extracted from −300 to 800ms relative to the events of interest. We compared six pairs of conditions: Global Go vs. Global Stop / Selective Stop / Ignore; Global Stop vs. Selective Stop / Ignore; and Selective Stop vs. Ignore, taking only into account trials with correct responses. Since successful Stop trials oversample the slower part of the Go-trial response time distribution, we quantified Global Go response time for each subject and selected only the slower half of Global Go trials for decoding. Trials were balanced in all condition pairs through random sampling of the condition with more trials. For decoding analysis, a linear Discriminant Analysis-based decoder was trained and tested at each time point using leave-one-out cross-validation, in which the decoder was trained on all but one trial and tested on the remaining trial until all trials were tested. Decoding performance was then quantified at each time point for each subject using the area under the curve (AUC) measure. The resulting 15 AUC time series data (551 timepoints) were compared against chance using paired samples t-tests, corrected for multiple comparisons using cluster permutation statistics (10,000 iterations, sample-wise p = .05; cluster-wise p = .01).

### Motoric Independent Component Selection

EEG signal components reflecting motor activity were identified from the ICA solution through an automated algorithm. Each independent component from each subjects’ were backprojected into channel space. The resulting activity at motor electrode sites (C3 and C4) was then converted into beta burst time series (see below). The resulting time series were then epoched with respect to the stimulus onset on correctly-responded Global go trials and then baseline-corrected with respect to the preceding 500ms period. The 200ms pre-response period for each trial was then saved and averaged across trials. The components with the largest reduction of beta bursting in this time period, relative to pre-stimulus baseline, were then selected as motor components, separately for left (C3) and right (C4) motor cortex.

### Time-frequency analyses

Full-spectrum event-related spectral perturbation (ERSP) plots (**Figure 3A**) were generated by transforming the EEG and LFP time series into the time-frequency domain via the filter-Hilbert method, with individual frequencies ranging from 1 to 125 Hz in linear steps of 1Hz and a band range of +/-.5Hz around each center frequency. The resulting complex time series was converted into power units through squaring its real part. Event-related time-frequency activity was then extracted from −500ms to 1,500ms around each event of interest and converted into change from baseline (decibel) using the 100ms prior to the event as a baseline. ERSPs were tested for significance via cluster-based nonparametric permutation testing (5,000 iterations, voxel-wise p = .05, cluster-wise p = .05) across the entire time-frequency spectrum.

Beta-frequency burst probability timelines (**Figure 3B**, **Figure 5C,D**) were extracted using established procedures (Shin et al., 2017, Enz et al., 2021; Diesburg et al., 2021). First, each electrode’s data were convolved with a complex Morlet wavelet of the form

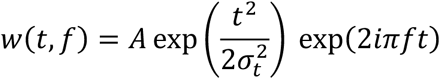

with 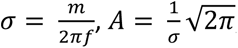, and *m* = 7 (cycles) for each of 18 linearly spaced frequencies spanning the β-band (13–30 Hz). Time-frequency power estimates were extracted by calculating the squared magnitude of the complex wavelet-convolved data. Individual β-bursts were defined as local maxima in the recording-wide, electrode-specific β-band time–frequency power matrix using the MATLAB function imregional(), using a set cut-off of 2× the median power in the case of EEG (Enz et al., 2021) and 6x the median power in the case of the local field potential (Shin et al., 2017; Diesburg et al., 2021). The resulting stick function was then converted into a continuous burst-rate timeline via convolution with a gaussian window of the form 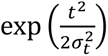, with a width of 200ms and *σ* = *m*/2*πf*, with m=7 (cycles) and being the highest frequency of the time-frequency composition (30Hz). The beta burst time series were tested for differences using sample-to-sample t-tests using the critical p-values indicated in the results section. Corrections for multiple comparisons were performed via the false discovery rate (FDR) method (Benjamini et al., 2006). Beta burst counts in the Stop/Ignore-signal-to-SSRT period (**Figure 3D**) were tested for significance using pairwise t-tests for each comparison.

### Connectivity analyses

For the purposes of all connectivity analyses involving the EEG data, the EEG data were transformed to current-source density (Tenke & Kayser, 2012) to reduce the impact of volume conduction.

The cortico-subthalamic connectivity plots in **Figure 5A** were generated by identifying beta bursts in each STN (using the methods indicated above) across each subject’s entire recording, and then identifying bursts in each channel’s EEG time series in the time window 50ms following each STN burst. The resulting averaged burst-rate scalp topography was then tested for differences from 0 at the group-level using one-tailed t-tests, corrected to a critical p-value of .00081 (i.e., p = .05 divided by the number of channels to correct for multiple comparisons).

The STN-to-STN connectivity plots in **Figure 5B** were generated by splicing the beta-burst rate time series (see above) for each STN across each subjects’ entire recording around individual bursts in the other STN, using a window of −1,000 to 1,000ms around each burst. The averaged time series were then tested against the mean burst probability across the entire recording (black line) for the respective STN using sample-to-sample t-tests, corrected for multiple comparisons using the FDR method.

The event-related STN-to-STN connectivity analyses in **Figure 5E** were performed by transforming the two STN time series into the frequency domain using the filter-Hilbert method as described above. Frequency-specific phase-angle differences were then extracted from the resulting complex time series, spliced around each event of interested, baseline-corrected to decibel as described above. They were then tested for significance via cluster-based nonparametric permutation testing (5,000 iterations, sample-wise p = .05, cluster-wise p = .05) across the entire time-frequency spectrum.

The SSRT-related cortico-subcortical connectivity analyses in **Figure 4A** were performed on correct Global-Stop, Ignore, and Selective-Stop trials by windowing the time-frequency power-series (extracted via the square of the real part of the filter-Hilbert transformed signal in the beta band, 13-30Hz) for each channel across four pre-SSRT windows of 50ms duration, starting from 200ms prior to SSRT, and across ten post-SSRT windows of 50ms duration, ranging from SSRT to 500ms after. Average channel-based beta power in each window was then correlated with the STN beta power in the 50ms pre-SSRT window used in **Figure 3D** (for the pre-SSRT windows) and with STN beta power in the 100ms post-SSRT window during which STN activity started diverging (**Figure 3B**) for the post-SSRT windows. Subject-specific SSRT estimates were used to time-lock the trial-wise data. The resulting correlation coefficients were then Fisher’s z-transformed and tested for significant differences from zero using channel- and window-wise t-tests to a critical p-value of p = .0063 for the pre-SSRT windows and .0025 for the post-SSRT windows (i.e., p = .05 divided by the number of time windows for each analysis).

The SSRT-locked, time-lagged, sample-to-sample, ROI-based cortico-subcortical connectivity analysis in **Figure 4B** was performed on correct Global-Stop, Ignore, and Selective-Stop trials by averaging the EEG signal within nine ROIs covering the entire scalp montage (ROIs indicated by the colored electrode locations), converting the nine resulting time-series to time-frequency power-series using the square of the real part of the filter-Hilbert transformed signal in the beta band (13-30Hz), and then extracting trial-wise activity around the subject-specific SSRT estimate. For each sample point in the −500ms to 1,500ms period around the subject-specific SSRT estimate, we then extracted and z-scored beta power in each EEG ROI and for each of the two STN power series, and then correlated the trial-wise beta power time series for each sample point. The resulting 1000×1000 temporal offset correlation matrix (2,000ms per trial at a sampling rate of 500Hz) was then converted to Fisher’s z and tested for significance via cluster-based nonparametric permutation testing (5,000 iterations), with a voxel-wise and cluster-wise critical p-value of .0028 (i.e., p = .05 divided by 18, the number of matrices across the 9 ROIs and 2 STN locations).

## Acknowledgments

This work was funded by NIH grant 2R01NS117753 to JRW, an endowment from the Clement T. and Sylvia H. Hanson Family to JRW, and a gift by Todd and Val Meyerhoff through the Kevin Dill Golf Tournament for Dementia, Parkinson’s and Veterans.

## Supplementary Materials

**Supplementary Table 1.** Clinical patient characteristics. Duration = Years since initial PD diagnosis. Hoehn-Yahr stage and Unified Parkinson’s disease rating scale (UPDRS) Subscale 3 scores quantified ON DBS. LEDD = Levodopa equivalent daily dose in milligrams. Exact Montreal Cognitive Assessment (MoCA) scores were not logged for the first five participants but were above 19.

| # | Duration<br>(years) | Hoehn-Yahr | UPDRS<br>III | LEDD<br>(mg/day) | MoCA |
| --- | --- | --- | --- | --- | --- |
| 1 | 20 | 3 | 29 | 813 | >19 |
| 2 | 22 | 4 | 33 | 625 | >19 |
| 3 | 13 | 2 | N/A | 1750 | >19 |
| 4 | 13 | 2 | 31 | 375 | >19 |
| 5 | 10 | 3 | N/A | 500 | >19 |
| 6 | 16 | 2 | 48 | 1225 | 24 |
| 7 | 10 | 2 | 24 | 250 | 21 |
| 8 | 8 | 2 | 15 | 660 | 23 |
| 9 | 8 | 2 | 33 | 731.25 | 25 |
| 10 | 6 | 2 | 45 | 750 | 25 |
| 11 | 4 | 3 | 35 | 375 | 26 |
| 12 | 7 | 2 | 53 | 2000 | 26 |
| 13 | 4 | 2 | 23 | 250 | 24 |
| 14 | 3 | 2 | 15 | 0 | 26 |
| 15 | 8 | 2 | 26 | 375 | 25 |
| 16 | 11 | 2 | 30 | 0 | 28 |

